# Early genomic detection of the Cosmopolitan Genotype of Dengue Virus 2 in Angola, 2018

**DOI:** 10.1101/424747

**Authors:** Sarah C Hill, Jocelyne Neto de Vasconcelos, Bernardo Gutierrez Granja, Julien Thézé, Domingos Jandondo, Zoraima Neto, Marinela Mirandela, Cruz dos Santos Sebastião, Ana Luísa Micolo Cândido, Carina Clemente, Sara Pereira da Silva, Túlio de Oliveira, Oliver G Pybus, Nuno R Faria, Joana Morais Afonso

## Abstract

We used portable genome sequencing to investigate reported dengue virus transmission in Angola. Our results reveal autochthonous transmission of dengue serotype 2 (Cosmopolitan genotype) in Jan 2018.

## Introduction

In Africa, the burden of disease caused by *Aedes-*borne virus infections may be similar to that of the Americas (1, 2). However, the transmission and genetic diversity of arthropod-borne viruses in Africa remains poorly understood due to a paucity of systematic surveillance. Moreover, syndromic surveillance may confound symptomatically similar illnesses {Grubaugh, 2018 #4149;Sharp, 2015 #4071} and current serological diagnostic tests can be masked by cross-reactivity to other circulating flaviviruses (3). Improved genomic surveillance of disease causing viruses can assist in better understanding the dynamics of transmission in Africa.

During 2013, Angola experienced a large dengue outbreak that was concentrated in Luanda Province (4). Cases detected in returning travellers showed that the virus rapidly disseminated from Angola to Europe, Asia and the Americas (5). Whilst infections were dominated by serotype 1 viruses (6), all dengue viral serotypes were reported in returning travellers from Angola (7).

Although dengue is likely endemic in Angola, patterns of DENV transmission in the country outside the 2013 epidemic are poorly characterized. The lack of genomic characterization restricts our understanding of DENV diversity within Angola, and the frequency and directionality with which dengue viruses are exchanged with other countries. Although enhanced national surveillance and genomic capacity can improve outbreak detection and response, this remains challenging in many public health laboratories due to the costs and laboratory capacity required for routine culture and bench-top sequencing (8). Recent technological advances now permit less expensive and pocket-sized sequencing using the MinION portable sequencer. Here we use a combination of portable sequencing and genetic analysis to detect the causative lineage of a DENV outbreak in Luanda.

### Epidemiological data

To investigate the timing and frequency of dengue occurrence in Angola, rapid diagnostic SD-Bio Line Dengue Duo (NS1 Ag+) tests were conducted to detect dengue specific IgM, IgG and NS1 presence. Samples were collected between 1st Jan 2016 and 15^th^ May 2018 from patients for which a physician suspected dengue as the cause of the patient’s illness presenting symptoms associated with DENV in central Luanda (n=6839, 3276 male). Samples were originally obtained for routine diagnostic purposes from persons visiting local clinics. In these cases, we used residual samples without informed consent with the ethical approval of the National Ethical Committee, Ministry of Health, Angola.

During Jan 2016 and mid May 2018, 80 NS1 positive cases were observed among a total of 6839 tested samples suspected for dengue infection. The first confirmed infections were detected in May 2017 and the number of cases seemed to peak around May 2018 (**Fig. 1A**).

**Figure 1.**
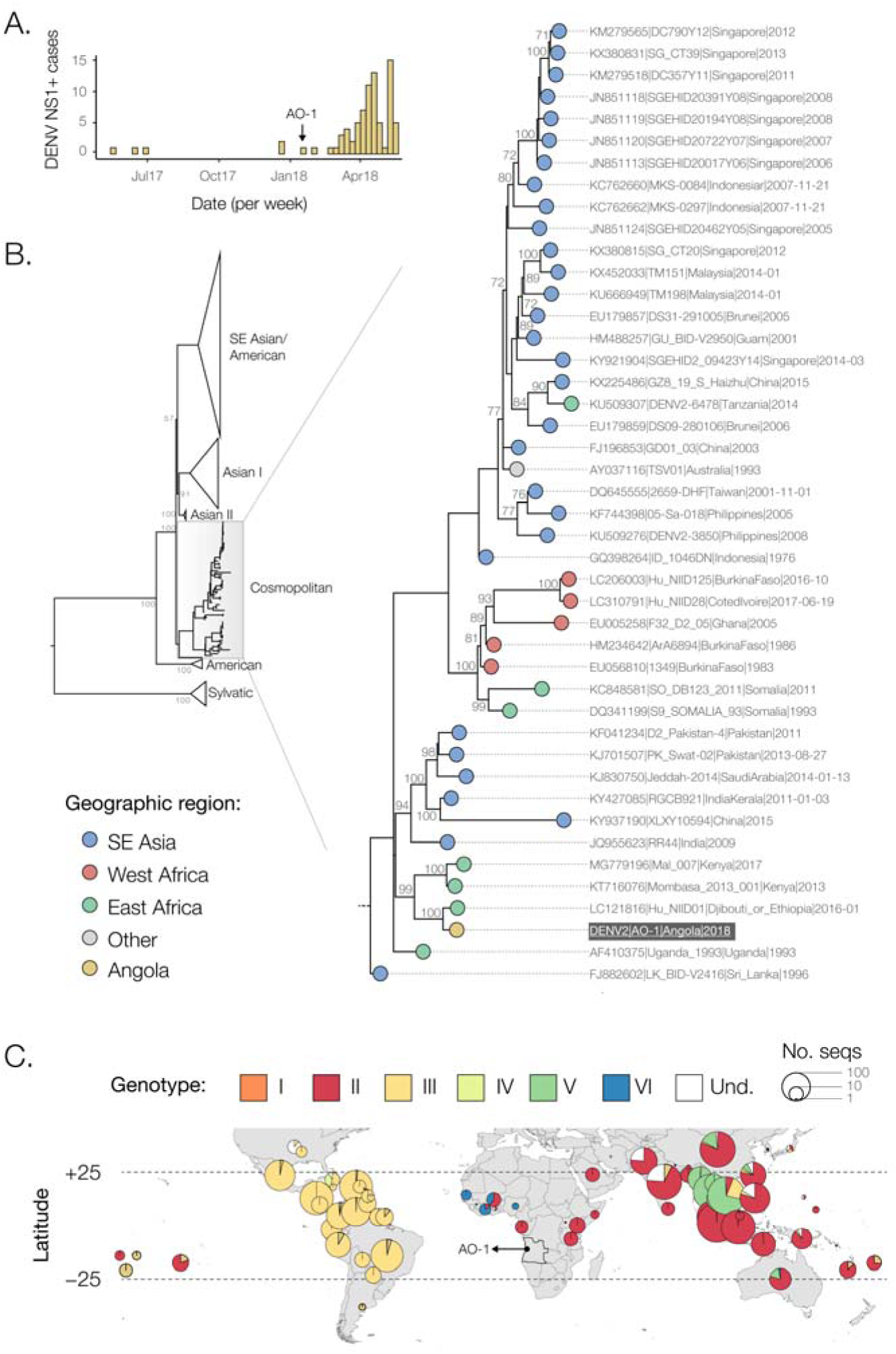
**A)** Number of dengue virus infections in Luanda (NS1 positive cases) from Jan 2016 to mid-May 2018. **B)** Midpoint rooted maximum likelihood phylogeny of DENV2 whole genomes. Support for branching structure is shown by bootstrap values at nodes. The Cosmopolitan genotype clade containing the Angolan DENV2 sequence is expanded. **On the right side,** an ML phylogeny of DENV2 Cosmopolitan genotype envelope gene sequences (1479bp), rooted on a sylvatic genotype outgroup that was included during phylogenetic tree estimation (not shown). Colors indicate geographic location of sampling. The Angolan DENV2 is highlighted. Support for branching structure is shown by bootstrap values at nodes (bootstrap scores >70 shown). **C)** Geographic distribution of availability of dengue serotype 2 sequence data (>100bp). Pie chart radii are log-proportional to the number of sequences available in each country, and are colored by genotype (und. = undefined).

### Molecular diagnostics

153 randomly selected sera samples (either IgG, IgM or NS1 positive) collected from 21^st^ July 2017 to 31^st^ Jan 2018 were tested for DENV RNA by real-time quantitative PCR (RT-qPCR) at the Instituto Nacional de Investigação em Saúde (INIS), Luanda, Angola using the CDC DENV1-4 Real-Time RT-PCR assay kit on an Applied Biosystems 7500 Fast machine, according to the manufacturer’s instructions (7). Of these, one sample, named herein as isolate AO-1, yielded a RT-qPCR cycle threshold of 22.5 for DENV-2. This sample was retrieved from a 48-yo male Luandan resident who visited a clinic on 18^th^ Jan 2018. The patient reported travelling to Mussulo Island between 24 Dec 2017 – 2 Jan 2018, a resort 30 km from Luanda.

### Nanopore genome sequencing

The RT-qPCR positive sample was subjected to viral genomic amplification and sequencing using a multiplex PCR primer scheme designed to amplify the entire coding region of DENV2 as previously described ^11^. Published genomes of non-sylvatic DENV2 were aligned and used to generate a 90% consensus sequence that formed the target for primer design. Primers that generate overlapping amplicons of length 980bp, with overlap 20bp were designed following (9) (**Table S1**). Details in cDNA synthesis, multiplex PCRs using 35 cycles, library preparation, sequencing, and generation of consensus sequences can be found in **Technical Appendix**.

The 90% consensus sequence defined above was originally used as a reference genome for mapping sequencing reads, but this reference was later refined to a more appropriate reference genome (Genbank Accession LC121816) using BLAST searching of the provisionally mapped data. The median sequencing depth was 11,448 reads, and 75% of the genome had a depth of at least 2419 reads. In total, 96% of the coding region of DENV2 was sequenced (named isolate AO-1, GenBank Accession number MH460898).

### Phylogenetic analyses

Phylogenetic trees were estimated to explore the relationship of the sequenced AO-1 genome to other isolates. 1395 DENV2 genome sequences with associated date and country of collection were retrieved from GenBank. From this data set, a subset was generated that included all identified African sequences (n=35), 200 globally sampled sequences (randomly sampled from the remaining 1360 sequences), and the novel AO-1 sequence. These sequences were aligned using MUSCLE, as implemented in Geneious 9.0.5 (10). A maximum likelihood (ML) phylogenetic tree was estimated using a GTR+4Γ substitution model in RAxML 8.2.10 (11). 500 non-parametric bootstrap replicates were performed to evaluate statistical support for phylogenetic nodes.

Most DENV sequences available in GenBank are partial gene sequences. We therefore supplemented the above whole genome alignment with all African DENV2 sequences >1000bp that belonged to the same genotype as the novel Angolan sequence. Sequences were aligned to the envelope region of DENV2, and a separate phylogeny was estimated from this alignment using the models specified above.

Phylogenetic estimation strongly supports placement of the Angolan genome in the Cosmopolitan genotype of DENV2 (**Fig. 1B**). The Angolan strain forms part of a well-supported monophyletic clade that comprises genomes sampled in East Africa, and is most closely related to DENV isolated from a returning traveler from this region. Inclusion of partial sequences in the phylogeny indicates that viruses from this clade have been present in East Africa since at least 2013 (**Fig. 1B**).

Finally, we generated maps of the distribution of currently available DENV sequence data, in order to explore the global distribution of the DENV2 Cosmopolitan genotype and to identify geographic gaps in DENV genomic surveillance that might bias phylogenetic interpretation. All DENV sequences >100bp of any serotype, with known location of sampling (including returning travelers), were downloaded from GenBank. DENV2 sequences were genotyped using the Genome Detective online classification tool (http://www.genomedetective.com). The majority of sequenced dengue viruses in Africa belong to DENV2 (49%), of which 70% belong to the Cosmopolitan genotype (**Fig. 1C**). We find that whilst 16% of all global clinically apparent dengue infections have been estimated to occur in Africa (2), DENV1-4 sequences from Africa currently represent <1% of the available global DENV sequence data. No data exists from the Democratic Republic of Congo, which has been epidemiologically linked with Angola during past arboviral outbreaks (12). Additional data will help to address transmission dynamics of DENV2 in the country and to identify common routes of virus importation into Angola.

## Conclusions

The DENV2 portable sequencing approach reported here represents a useful tool for early genomic characterization and molecular epidemiology of DENV2 outbreaks in Africa and elsewhere. Based on phylogenetic evidence and the geographic distribution of detected genotypes, the DENV2 Cosmopolitan genotype detected in Angola is likely endemic in Africa. The AO-1 genome analyzed here probably represents an early transmission event from an ongoing DENV2 epidemic in Luanda. Further sequencing of DENV in the region is required to determine whether the Cosmopolitan genotype is endemic to Angola, or if it represents a more recent introduction from elsewhere (for example, from East Africa or from other unsampled locations).

## Acknowledgements

This study was made possible by funding from the Wellcome Trust and Royal Society Sir Henry Dale Fellowship (grant 204311/Z/16/Z), internal HEFCE Global Challenges Research Fund grant 005073, John Fell Research Fund Grant 005166). Travel to Angola by SCH and NRF was supported by Africa-Oxford Travel Grants (grant numbers AfiOx-48 and AfiOx-60). We would like to thank to all the patients involved and the staff who assisted with sample collection. This work forms part of the ArboSPREAD project.

## Technical Appendix

### DENV2 complete genome MinION nanopore sequencing

Between the 15 and 23^rd^ February 2018, we attempted sequencing using the Oxford Nanopore MinION device at Instituto Nacional Investigação em Saúde (INIS), Ministry of Health of Angola, Luanda, on the AO-1 isolates, as part of the ArboSPREAD project focused on genomic surveillance of arthropod-borne viruses. Diagnostic, sequencing and genetic analysis results were presented at the INIS to local public health authorities on the 23^rd^ February 2018. We used the protocol chemistry R9.4, and the detailed protocol has been previously described in (9). Briefly, the protocol starts with cDNA synthesis using random primers and is followed by gene-specific multiplex PCR (9). Extracted RNA was converted to cDNA using the Protoscript II First Strand cDNA synthesis Kit (New England Biolabs, Hitchin, UK) and random hexamer priming. DENV2 genome amplification by multiplex PCR was attempted using the primer scheme shown in **Table S1** and 35 cycles of PCR using Q5 High-Fidelity DNA polymerase (NEB). PCR products were cleaned up using AmpureXP purification beads (Beckman Coulter, High Wycombe, UK) and quantified with the Qubit dsDNA High Sensitivity assay on a Qubit 3.0 instrument (Life Technologies). PCR products for the AO-1 sample yielded sufficient material and were thus barcoded and pooled in an equimolar fashion using the Native Barcoding Kit (NBD103, Oxford Nanopore Technologies, Oxford, UK). Sequencing libraries were generated from the barcoded products using the Genomic DNA Sequencing Kit SQK-LSK208 (Oxford Nanopore Technologies). We used 250 ng of total DNA input in the library preparation. The library was loaded onto a R9/R9.4 flow cell (FLO-MIN106) and sequencing data were collected for up to 48□hr. Consensus genome sequences were produced by alignment of two-direction reads first to a 90% consensus sequence, then to a DENV2 reference genome (GenBank Accession number: LC121816). Positions with□≥□20× genome coverage were used to produce consensus alleles, while regions with lower coverage, and those in primer-binding regions were masked with N characters.

**Table S1.**
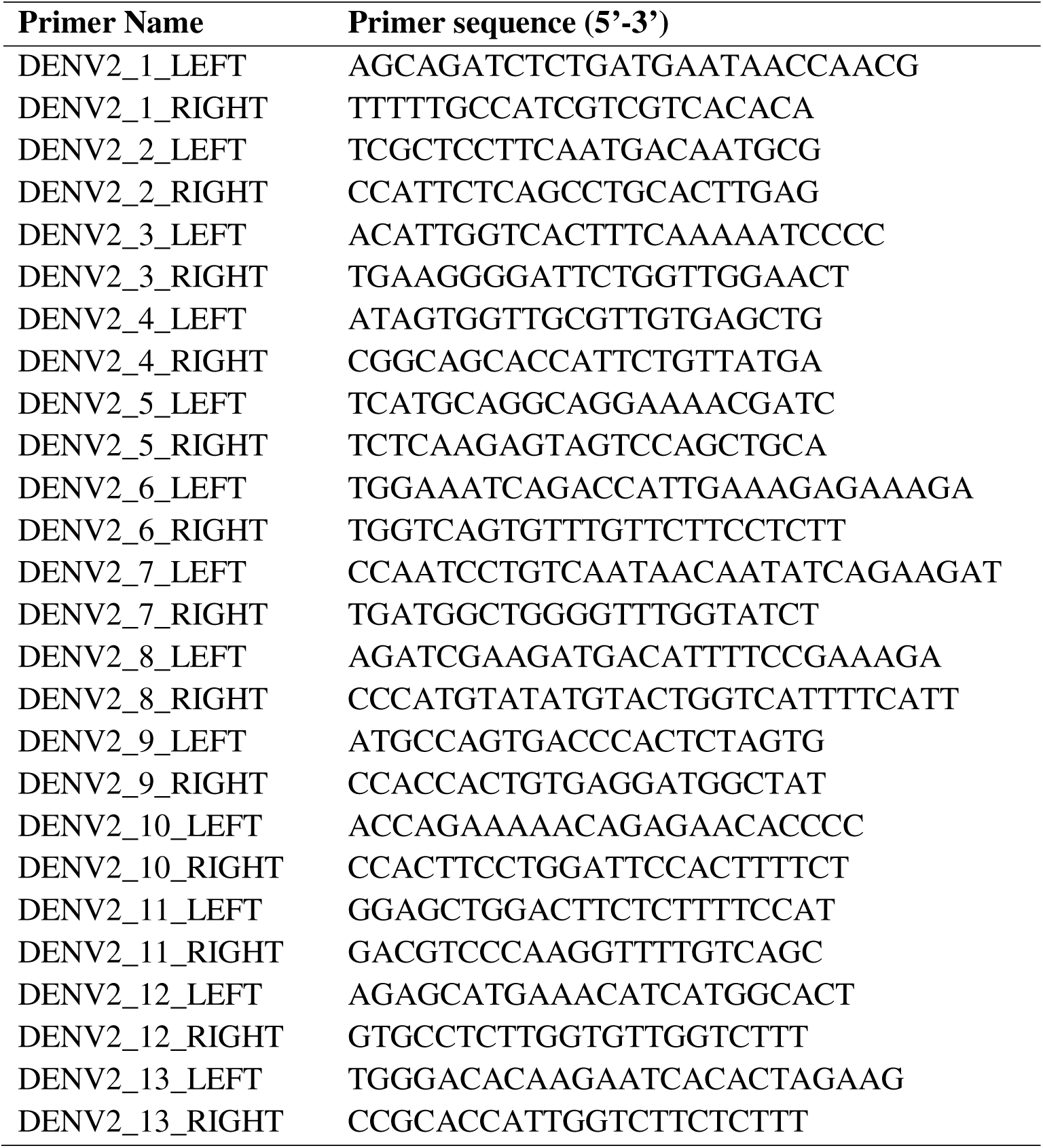
Sequencing primers used for MinION sequencing of DENV2.

**Figure S1.**
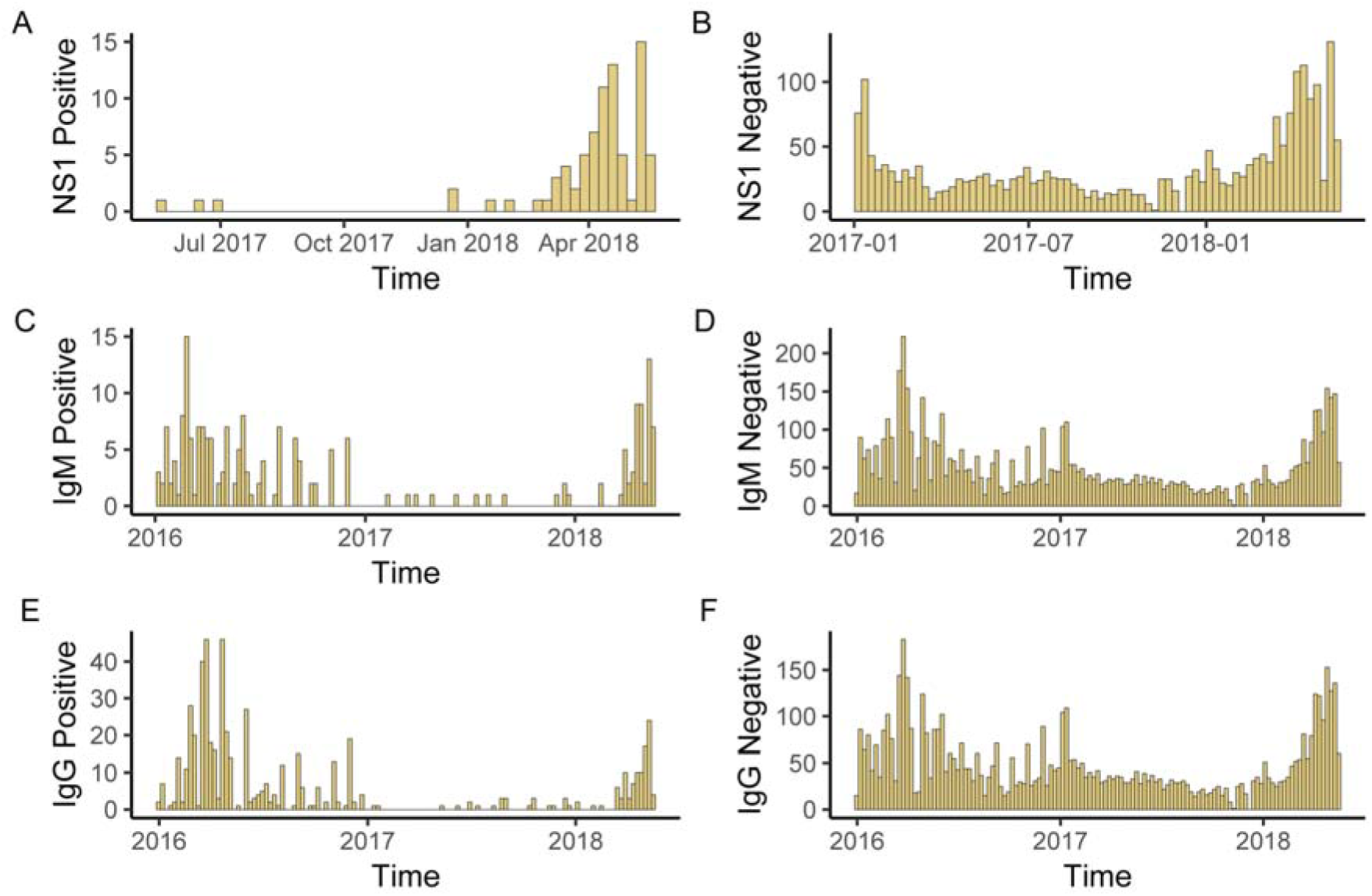
Summary of the serological results obtained from 6839 tests performed in Luanda between Jan 2016 and 15^th^ May 2018. Panels A, C and E show positive results; panels B, D and F indicate number of negative results through time. Note that while IgM (panels C and D) and IgG (panels E and F) screening started in Jan 2016, NS1 screening (panels A and B) started only in Jan 2017. The IgM and IgG positive cases throughout 2016 are possibly due to the presence of antibodies against yellow fever virus.

